# Savings in visuomotor learning is associated with connectivity changes within a cerebello-thalamo-cortical network encoding movement errors

**DOI:** 10.1101/2024.09.13.610754

**Authors:** Lucas Struber, Laurent Lamalle, Pierre-Alain Barraud, Aurélien Courvoisier, Rafaël Laboissière, Takayuki Ito, Vincent Nougier, David J Ostry, Fabien Cignetti

## Abstract

Savings refers to faster relearning upon re-exposure to a previously experienced movement perturbation. One theory suggests that the brain recognizes past errors and is therefore more able to learn from them. If true, there should be a modification of the neural response to errors during re-exposure to a perturbation. To test this idea, we imaged the brains of participants who underwent two sessions (1 day apart) of adaptation to a visuomotor perturbation and investigated brain responses to movement errors. The magnitude of movement error was entered into different types of GLMs to study error-related activation and coactivation (or functional connectivity). We identified a cerebello-thalamo-cortical network involved in the processing of movement errors during adaptation. We found that connectivity between regions of this network (i.e., between the cerebellum and the thalamus, and between the primary somatosensory cortex and the anterior cingulate cortex) became stronger during re-adaptation. Importantly, participants with the largest increases in connectivity strength were those who demonstrated the largest amounts of savings. These results establish a relationship between the ability of the brain to represent errors and the phenomenon of savings.

## Introduction

The formation of motor memory has typically been studied in the context of adaptation paradigms in which subjects learn to compensate for a systematic perturbation, mostly involving a manipulation of the visual feedback (Krakauer, 2009; Krakauer et al., 2005) or a change in the dynamics of the motor apparatus (Shadmehr and Brashers-Krug, 1997; Shadmehr and Mussa-Ivaldi, 1994). In these paradigms, one important observation is that subjects learn to compensate for the perturbation in fewer trials when they have previously experienced that perturbation, a phenomenon referred to as savings (Zarahn et al., 2008; Smith et al., 2006; Krakauer et al., 2005; Kojima et al., 2004). Savings during repeated perturbation exposures has been commonly assumed to result from the reemergence of a latent motor memory (Huang et al., 2011; Krakauer and Shadmehr, 2006). In particular, several studies have reported that when subjects first encounter a perturbation, they learn to adjust their behavior to compensate for the perturbation, such as aiming in the opposite direction of the rotation of the visual feedback during visuomotor adaptation (Taylor et al., 2014; Taylor and Ivry, 2011). Later, upon re-exposure to the same perturbation, individuals recall this action, resulting in faster relearning (Avraham et al., 2021; Morehead et al., 2015). Alternatively, converging lines of evidence have established that savings does not necessarily result from the recall of a previous action but occurs through an increase in the sensitivity to movement errors during relearning (Coltman et al., 2019; Leow et al., 2016; Herzfeld et al., 2014b). Experiencing a particular error at one time leads to a durable change in response to the same error in the future, the brain becoming more able to learn from that error. This view is reinforced by studies having shown that the brain can tune its rate of learning based on previous error history (Coltman et al., 2021; Gonzalez Castro et al., 2014; Tan et al., 2014; Braun et al., 2009; Burge et al., 2008). Thus, savings appears to depend not only on a memory of actions but also on a memory of errors.

At the brain level, there is evidence that memories formed during motor adaptation are stored in cortical motor and somatosensory regions (Ebrahimi and Ostry, 2024; Galea et al., 2011; Landi et al., 2011; Hadipour-Niktarash et al., 2007). However, it remains unclear whether these regions genuinely support faster relearning. Modulation of the excitability of these regions using transcranial magnetic stimulation has sometimes been reported to affect savings (Kumar et al., 2019; Villalta et al., 2015), but sometimes not (Darainy et al., 2023). Similarly, there is no general consensus among functional neuroimaging studies regarding which brain re-gions and mechanisms participate in savings. Some studies failed to find any relationship between brain activity and savings (Della-Maggiore et al., 2017; Bédard and Sanes, 2014), whereas others reported associations, but mainly with non-sensorimotor regions such as the hippocampus and default-mode areas (Standage et al., 2023; Gale et al., 2022; Cassady et al., 2018). However, changes in the neuronal population activity dynamics of the monkey primary motor cortex has been found to correlate with faster relearning (Sun et al., 2022). Furthermore, there is some evidence that the cerebellum may also play a role in the retention of motor memory (Herzfeld et al., 2014a), including savings seen during relearning (Medina et al., 2001). In particular, an fMRI study demonstrated increased activity in the cerebellum (lobule VI) when repeating a motor adaptation task, which correlated with the amount of savings (Debas et al., 2010). Finally, in locomotor adaptation, a relationship was found between cerebellar-thalamic intrinsic functional connectivity and faster relearning (Mawase et al., 2017). Therefore, savings might be supported by neural changes that take place in a large scale cerebello-thalamo-cortical network.

Here, we assessed the neural basis of savings by asking whether this phenomenon builds specifically on brain areas coding for errors. It has long been established that the brain is tuned to errors, including the sensorimotor cortical and cerebellar regions as well as the parietal association areas during motor performance (Luauté et al., 2009; Grafton et al., 2008; Diedrichsen et al., 2005). Importantly, these regions not only serve as a passive observer that encodes the errors, but they also play an active role in correcting these errors. Electrical stimulation delivered to these areas evokes movements that compensate for the errors (Inoue and Kitazawa, 2018; Inoue et al., 2016). Furthermore, changes in intrinsic functional connectivity following learning are predominantly found in error-related regions (Bernardi et al., 2018), suggesting that they are key regions for motor memory formation. These properties suggest that when we become better at learning, manifesting savings, it is because error-related brain regions have undergone changes. Specifically, we hypothesized that these changes may apply to activation and coactivation (i.e., functional connectivity) properties of the error-related brain areas. To test this assumption, we conducted an fMRI visuomotor learning/relearning (24h later) experiment and assessed what changes in activation and functional connectivity of brain regions that processed errors (as identified using a parametric modulation approach) accompanied savings. We also looked for changes in intrinsic functional connectivity following learning and relearning sessions to get insights into brain changes that may have survived beyond the task.

## Methods

### Population

Twenty-four healthy right-handed young adults (age: 29.4 ± 3.6 years old; 11 women) participated in this study. They were free of any neurological or musculoskeletal injuries, and presented normal or corrected-to-normal vision. The study was conducted with the approval of the ethics committee “Comité de Protection des Personnes Nord Ouest IV” under the approval ID-RCB n° 2020-A00268-31. Written informed consent was obtained from all participants.

An a priori power analysis was conducted using G*Power 3.1.9.7 for sample size estimation (Faul et al., 2007), based on data from (Standage et al., 2020). This study investigated visuomotor adaptation on two testing days in a sample of 32 subjects, assessing whether the rate of learning increased over the two days (savings). The effect size was d=1.08 - obtained from t-value using the formula 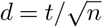 (Rosenthal, 1991) – which can be interpreted as a large effect size (Cohen, 1988). With a significance criterion of α = .05 and power = .80, the minimum sample size needed with this effect size was n = 7 for a paired t-test. Furthermore, Standage et al. (2020) identified distinct profiles of learners across days, including one profile with poor savings. Using data of this subsample (n=10) and using the same parameterization as above, the sample size needed was n=24. Thus, our sample size of n = 24 should be adequate to reveal savings, even in the least favorable scenario.

### Experimental setup

The experiment took place on two consecutive days, which both included an anatomical scan, resting-state runs, and task-based runs (Figure 1). To reduce head motion and scanner noise, foam padding and earplugs were provided to the participants. During anatomical scans, participants were instructed to lie quietly looking at a black cross, which was displayed on the screen with a grey background to avoid visual fatigue. For the task-based runs, participants held with their right hand a custom-made fMRI-compatible joystick. A Plexiglas adjustable table fixed above their pelvis enabled them to comfortably rest their arm and control the joystick while minimizing movements of the elbow. Participants were instructed to perform the task with their forearm trying to avoid as much as possible large arm movements. Visual stimuli were back-projected onto a monitor (60-Hz frame rate, 3840 × 2160 pixels screen resolution, 40” diagonal, NordicNeuroLab®) and viewed through a mirror mounted on the head coil. The task consisted in performing target aiming movements with the joystick that controlled a green cursor (Struber et al., 2021). Each movement started from the center of the screen to one of eight possible targets equally spaced around a virtual circle (radius 40 cm). Each trial started with one of the eight targets becoming red. Participants had to reach the target with the green cursor as fast and as accurately as possible. Once the target was reached, it turned to blue and participants had to maintain the cursor inside the target until the target disappeared and a red circle appeared in the middle of the screen indicating them to passively let the joystick return to its initial position (Figure 1). Allotted time was 2 s for target reaching and 1.45 s for passive return movement. These specifications allowed us to collect fMRI data during 36 repetition times (TR) per block. The task was implemented using a custom C++ software based on Qt and Measurement Computing© libraries. This software was synchronized with the MRI scanner.

**Figure 1.**
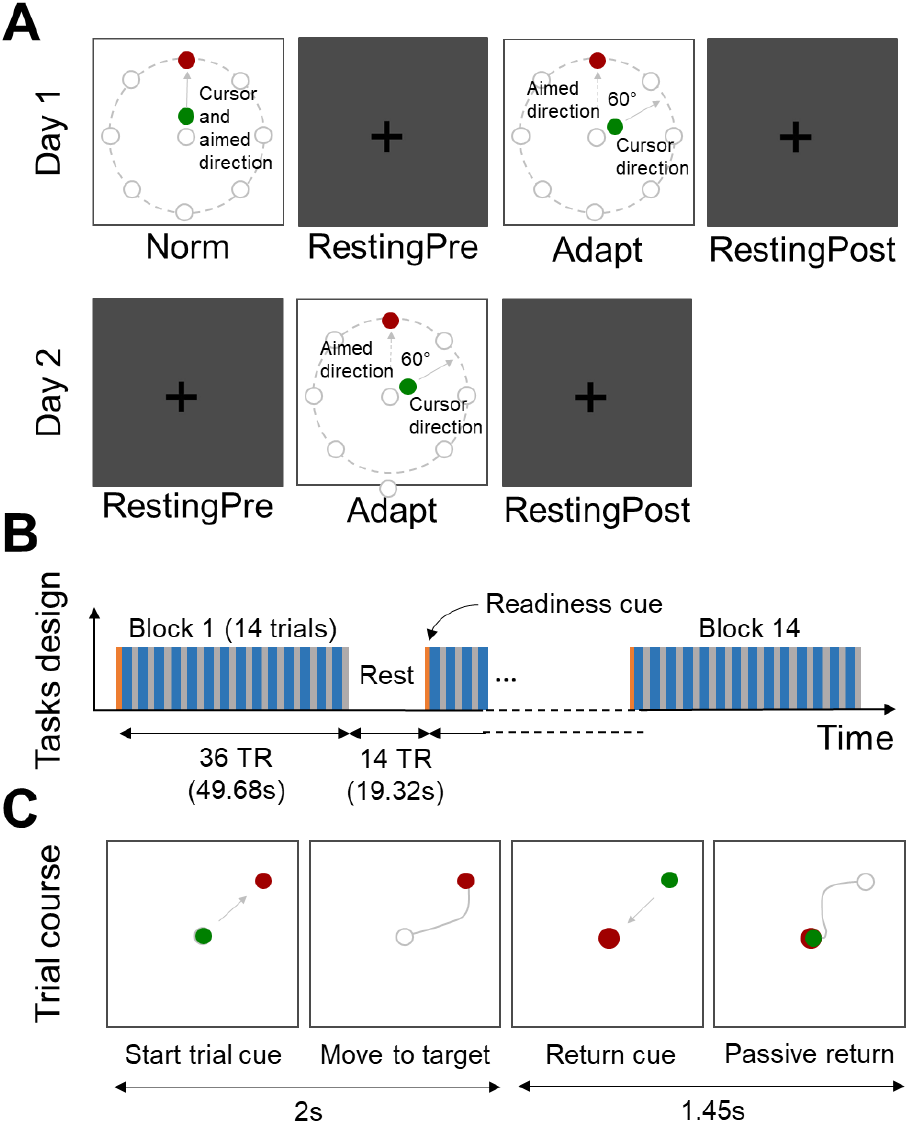
Design of the experiment. **(A)** Sequence of experimental tasks during the two consecutive days of the experiment. Day1 started with the Norm condition, in which the cursor direction was consistent with participants’ aimed direction. It was followed by a pre-adaptation resting-state, then the adaptation condition, in which the cursor direction was 60°clockwise rotated with respect to the aimed direction. Finally, another resting-state was performed postadaptation. In Day2, the same sequence was used except the norm condition of Day 1. (**B**) Block design and timing of behavioral tasks (Norm, Adapt-day1 and Adapt-day2) in which 14 blocks of 14 trials were performed. Each block lasted 36 TR (MRI repetition time = 1.38 s), and were intertwined by a resting period of 14 TR. (**C**) Cues that were seen by participants for each trial, which was divided in two phases, the reaching phase during which the target was displayed in red and the cursor in green, and the passive return phase during which the center of the screen was colored in red. Grey lines are drawn only for comprehension purposes and were not visible by participants.

### Experimental conditions

Participants had to adapt to a constant perturbation on two consecutive days (Adapt-day1 and Adapt-day2). A 60°clockwise (CW) rotation angle was maintained between the cursor on the screen and the actual movement performed by the participants. We deliberately chose to not include null trials (i.e., washout) between the two days of adaptation to avoid unwanted anterograde interference and magnify, as much as possible, the extent of savings (Villalta et al., 2015). On day1, a normal, unrotated, movement condition was included before the adaptation condition so as to identify seed regions afterwards used for connectivity analysis (cf. fMRI Processing). Resting-state scans were also acquired on both days immediately before and after the adaptation task (RestingPre-day1, RestingPost-day1, RestingPre-day2 and RestingPost-day2). This was included so as to investigate possible changes in intrinsic connectivity following adaptation. Each task condition included 14 blocks of 49.68 s (36 TR) which started with 1 TR of readiness indicating the beginning of the block, followed by 14 trials of aiming movement with the cursor for a total of 196 trials per condition. Each block was followed by 19.32 s (14 TR) of rest (Figure 1).

### Behavioral analysis and modelling

Cursor positions acquired at 2048 Hz were filtered through a dual 4th order low pass Butterworth filter with a 5 Hz cutoff frequency and were then down-sampled at 100 Hz. Beginning of reaching movement was defined as the time the cursor velocity was above 20% of peak velocity. End of reaching movements was defined as the time the cursor entered into the target. Trials in which participants did not reach the target were excluded from further analyses (1.5%, 6.1% and 4.9% of the trials in Norm, Adapt-day1 and Adapt-day2 conditions, respectively). For each trial, the motor performance was quantified by the normalized cumulative error performed by the participant during target reaching. To do so, at each time step, the error was defined as the distance between the actual cursor position and the shortest trajectory (i.e., the straight-line connecting the starting position and the target). Then, all errors were summed (square root of the sum of squared distances) and normalized by reaching duration. This variable was used afterwards in parametric modulation of brain activation (see below). For each participant, learning was studied by modeling error over trials through a statespace model (SSM) (Diedrichsen et al., 2005; Donchin et al., 2003). Specifically, we modeled learning on each day separately (i.e. for each Adapt condition), using a one-state SSM of the following form (Albert and Shadmehr, 2018):

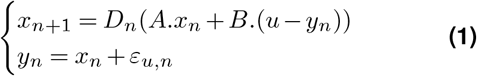

where

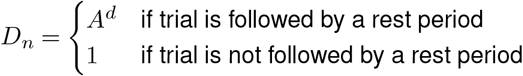

with *x*_*n*_ the hidden state at trial *n, u* the external perturbation, *y*_*n*_ the motor output, *A* the retention factor that controls the decay rate of the state in the absence of error due to the passage of time, *B* the error sensitivity parameter that controls the learning rate of the state from the error, *D*_*n*_ a parameter that depends on the decay factor *d* and serves to model the additional decay that elapses after the conclusion of the trial preceding a break (every 14 trials), and *ε*_*u,n*_ the motor noise representing the alteration of the generated movement with respect to the desired one. *x*_0_, the initial state of the model, encodes the priors of the learner. For example, a completely naïve learner would start with *x*_0_ = 0. *A, B, x*_0_ and *d* were parameters to be determined by optimization. To do so, the mean square error between actual error and prediction error *u*−*y*_*n*_ was minimized (using Matlab fmincon function and interior-point algorithm with optimality and constraint tolerances set to 10*−*15) to obtain the optimal values of parameters (LMSE algorithm). Our constrained search parameter space was defined by its lower and upper bounds: *A*∈ [0.1; 1], *B*∈ [0.1; 1], *x*_0_ ∈ [0; 2*u*] and *d*∈ [0.1; 30] (Albert and Shadmehr, 2018). We also derived an extra parameter from the model equation, namely the asymptotic value of motor error (Albert et al., 2021):

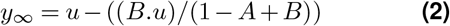

which informed on learning saturation following prolonged exposure to the perturbation. Differences in model parameters between days 1 and 2 were assessed using paired t-tests with a level of significance set at *p* < 0.05. The amount of savings was quantified as the difference in *B*-values between days 1 and 2, any increase of *B* reflecting savings. Analysis and statistical testing were performed using Matlab (R2018b).

### MRI acquisition

MRI data were acquired at 3T (Achieva dStream 3.0T TX, Philips, NL) with a 32-channel head coil at IR-MaGe MRI facility (Grenoble, France). To account for different head placements into the MRI, structural T1-weighted images were acquired at the beginning of each day, using a Magnetization-Prepared Rapid Acquisition Gradient Echoes (MPRAGE) (TI = 900 ms, TR/TE = 8.2 ms/4.7 ms, 220 slices, in-plane resolution = 1 × 1 mm, slice thickness = 1 mm, flip angle = 8°, field of view = 256 × 256 × 220 mm^3^). Compressed SENSE with acceleration factor 4.4 was used. Resting scans functional blood-oxygen level-dependent (BOLD) images were collected using a T2*-echo-planar sequence with multiband acceleration of 3 and SENSE factor of 2 (TR/TE = 1620/30 ms, voxel size = 2.25 × 2.25 × 2 mm^3^, gap = 0.25 mm, 69 slices, field of view = 216 × 216 × 155 mm^3^, flip angle = 70°, 350 volumes). Task-based functional BOLD images were obtained with a T2*-echo-planar sequence with multiband acceleration of 3 and SENSE factor of 2 (multiband factor = 3, TR/TE = 1380/30 ms, voxel size = 2.5 × 2.5 × 2.25 mm^3^, gap = 0.25 mm, 63 slices, field of view = 200 × 218 × 157 mm^3^, flip angle = 70°, 700 volumes). To correct for magnetic field inhomogeneity during data preprocessing, we also acquired a pair of spin-echo images before each BOLD scan (same specifications as above for task-based or resting), with reversed phase encoding direction.

### MRI preprocessing

After conversion into a BIDS dataset (Gorgolewski et al., 2016), preprocessing was performed using FMRIPREP version 20.2.6 (Esteban et al., 2019, RRID:SCR_016216), a Nipype (Gorgolewski et al., 2016, 2011, RRID:SCR_002502) based tool. Each T1w (T1-weighted) volume was corrected for INU (intensity non-uniformity) using N4BiasFieldCorrection v2.1.0 (Tustison et al., 2010) and skull-stripped using antsBrainExtraction.sh v2.1.0 (using the OASIS template). Brain surfaces were reconstructed using recon-all from FreeSurfer v6.0.1 (Dale et al., 1999, RRID:SCR_001847), and the brain mask estimated previously was refined with a custom variation of the method to reconcile ANTs-derived and FreeSurfer-derived segmentations of the cortical gray-matter of Mindboggle (Klein et al., 2017, RRID:SCR_002438). Spatial normalization to the ICBM 152 Nonlinear Asymmetrical template version 2009c (Fonov et al., 2009, RRID:SCR_008796) was performed through nonlinear registration with the antsRegistration tool of ANTs v2.1.0 (Avants et al., 2008, RRID:SCR_004757), using brain-extracted versions of both T1w volume and template. Brain tissue segmentation of cerebrospinal fluid (CSF), white-matter (WM) and gray-matter (GM) was performed on the brain-extracted T1w using fast (Zhang et al., 2001, FSL v5.0.9, RRID:SCR_002823).

Functional data was slice time corrected using 3dTshift from AFNI v16.2.07 (Cox, 1996, RRID:SCR_005927) and motion corrected using mcflirt (Jenkinson et al., 2002, FSL v5.0.9,). Distortion correction was performed using an implementation of the TOPUP technique (Andersson et al., 2003) using 3dQwarp (AFNI v16.2.07). This was followed by co-registration to the corresponding T1w using boundary-based registration (Greve and Fischl, 2009) with nine degrees of freedom, using bbregister (FreeSurfer v6.0.1). Motion correcting transformations, field distortion correcting warp, BOLD-to-T1w transformation and T1w-to-template (MNI) warp were concatenated and applied in a single step using antsApplyTransforms (ANTs v2.1.0) using Lanczos interpolation. ICA-based Automatic Removal of Motion Artifacts (Pruim et al., 2015, ICA-AROMA) was performed on the preprocessed functional data after spatial smoothing with an isotropic, Gaussian kernel of 6mm FWHM (full-width half-maximum). Noise components were partially regressed in the internal regression step of ICA-AROMA, which constitutes an effective strategy to abolish motion-related variance for relatively low cost in terms of data loss (Parkes et al., 2018; Ciric et al., 2017). Additionally, region-wise global signals within WM and CSF were computed. Many internal operations of FMRIPREP use Nilearn (Abraham et al., 2014, RRID:SCR_001362), principally within the BOLD-processing workflow. For more details of the pipeline see https://fmriprep.readthedocs.io/en/20.2.6/workflows.html.

### fMRI Processing

Task-based and resting-state (participant-level and group-level) analyses were performed using custom-made (batch syntax) Matlab routines (R2018b) running SPM12 toolbox (https://www.fil.ion.ucl.ac.uk/spm/) and CONN toolbox (https://web.conn-toolbox.org/), respectively.

#### GLM analysis

Brain activation and functional connectivity during the adaptation sessions were assessed using standard and beta series (least squares-all method) GLMs, respectively (Rissman et al., 2004). Beta series GLM differ from standard GLM in that the model is fitted to the fMRI data trial-wise, and not condition-wise. This creates time series of beta estimates which can be subsequently correlated to estimate functional connectivity. Here, the GLM design matrix included movement trials, each trial being represented as a boxcar function timelocked to the onset of movement and convolved with SPM canonical hemodynamic response function (HRF). Trials were represented all together in a single column of the design matrix in standard GLM while they were represented separately in as many columns as there were trials in beta series GLM. A parametric modulator was also entered into the design matrix to model the linear effect relating the evoked-response during each trial to the magnitude of movement error. This modulator was entered the same way as the movement trials in the GLM, i.e., in a single column for standard GLM and in as many columns as there were trials in beta series GLM. This parametric modulator was demeaned to avoid collinearity issues prior to convolution with the HRF (Mumford et al., 2015). The WM and CSF time series were further entered into the design matrix to serve as nuisance variables. As for standard GLM, individual beta maps of the parametric modulator were then entered into a second-level analysis. One-sample t-tests revealed brain regions whose activation related to movement errors during the adaptation sessions. A two-sample t-test assessed whether activation of these regions related to errors was different between adaptation and re-adaptation sessions. As for beta series GLM, we extracted the mean beta series of the parametric modulator within four regions of interest (ROIs), including the primary motor cortex (M1), the primary somatosensory cortex (S1), and the lobules VI and VIIIb of the cerebellum. These lobules of the cerebellum are known to assume sensorimotor function (King et al., 2019; Buckner et al., 2011). As mentioned in introduction, these regions appear to be key regions for the storage of motor memories. ROIs were 6mm radius spheres built after running a standard GLM on the Norm condition and extracting coordinates of local maxima of the group-level (one-sample t-test) activation map (Norm versus implicit baseline). ROIs were centered at MNI coordinates [-38, -18, 62] for left M1, [-34, -30, 52] for left S1, [28 -48 -26] for right cerebellar lobule VI, and [18 -60 -54] for right cerebellar lobule VIIIb (Supplementary Figure S1). The fact that ROIs were obtained from a condition (i.e., Norm) other than the condition under study (i.e., Adapt) was done to contain the risk of circular analysis (Kriegeskorte et al., 2009). As a second step, the beta series of the parametric modulator of each ROI was correlated (Pearson) with the beta series of the parametric modulator of each voxel in the rest of the brain. We used the doBetaSeries function from Andy’s brain blog (http://andysbrainblog.blogspot.com/2014/06/beta-series-analysis-in-spm.html). This yielded correlation maps that were finally converted to z-score maps and entered into the second-level analysis. One-sample t-tests revealed co-activation, i.e. functional connectivity, patterns of the ROIs during the adaptation sessions. It is worth mentioning, given the methodology described above, that these patterns represented similarities between ROIs and brain regions in their responses to error during the task. Finally, two-sample t-tests assessed whether the co-activation patterns of the ROIs were different between adaptation and re-adaptation sessions.

#### Resting-state connectivity

Functional connectivity before and after adaptation sessions was scrutinized using seed-to-voxels and independent component (ICA) approaches. Seed-to-voxels maps were computed as the Fisher-transformed bivariate correlation coefficients between denoised average BOLD time series computed across all the voxels within each ROI presented above and each individual voxel of denoised BOLD time series in the brain. Group-level ICA followed Calhoun’s general methodology (Calhoun et al., 2001) and estimated 40 temporally coherent networks from the resting-state fMRI data combined across all subjects and conditions. The BOLD signal from every timepoint and voxel in the brain was concatenated across subjects and conditions along the temporal dimension. A singular value decomposition of the z-score normalized BOLD signal (subject-level SVD) with 64 components separately for each subject and condition was used as a subject-specific dimensionality reduction step. The dimensionality of the concatenated data was further reduced using a singular value decomposition (group-level SVD) with 40 components, and a fast-ICA fixed-point algorithm with hyperbolic tangent (G1) contrast function was used to identify spatially independent group-level networks from the resulting components. GICA3 back-projection was then used to compute ICA maps associated with these same networks separately for each individual subject and condition. Finally, we identified the network we were interested in, namely the sensorimotor network, using the correlational spatial match-to-template tool of CONN. Any difference that may have occurred between the resting state conditions were examined from F-tests run on seed-to-voxel and ICA (sensorimotor) maps.

#### Thresholding of statistical maps

Maps were thresholded using a cluster-forming threshold *p* < 0.001 and a cluster-extent based threshold *p* < 0.05 family-wise error rate (FWER) corrected, according to established recommendations (Eklund et al., 2016; Woo et al., 2014). Cluster-extent threshold was estimated using the Gaussian random field method (Worsley et al., 1996), as implemented in SPM 12 and CONN 19.

## Results

### State space modelling of behavior

The evolution of error across trials and model prediction on days 1 and 2 are plotted on Figure 2A (averaged across participants). Paired t-tests run on SSM parameters (Figure 2B) showed that the error sensitivity that controls the rate at which the state learns from previous error was significantly higher in day2 than day1 (mean ± SE: 0.052 ± 0.013 in day1, 0.106 ± 0.020 in day2, *p* = 0.036) while the retention factor that encodes the rate at which state decays with the passage of time remained similar whatever the day (mean ± SE: 0.995 ± 0.001 in day1, 0.992 ± 0.001 in day2, *p* = 0.186). Hence, there was a larger amount of learning from error during relearning, consistent with the notion of savings, while the amount of forgetting between trials was unchanged. We also observed that relearning saturated at a lower asymptotic error in day2 than day1 (mean ± SE: 0.122 ± 0.010 in day1, 0.089 ± 0.007 in day2, *p* = 0.007). This was a direct consequence of the fact that the amount of learning increased from day1 to day2 while the amount of forgetting remained the same (cf. equation 2). Finally, there was no change in the initial error (mean ± SE: 0.362 ± 0.017 in day1, 0.342 ± 0.014 in day2, *p* = 0.327) from day1 to day2, indicating that learning and relearning both started from the same value of internal state and testified that savings was unbiased.

**Figure 2.**
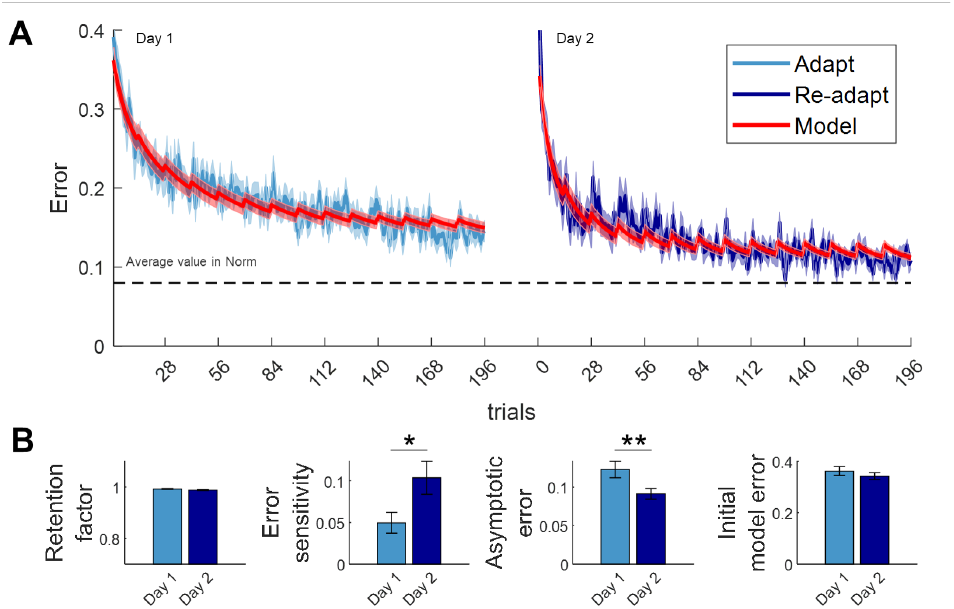
Behavioral and computational results. (**A**) Evolution of movement error across trials during Adapt-day1 (light blue) and Adapt-day2 (dark blue) conditions, and as predicted by the state space model (internal state in red). (**B**) Bar graphs of the model parameters, with retention factor = *A*, error sensitivity = *B*, and initial model error = *x*_0_ in equation 1, and asymptotic error = *y*_*∞*_ in equation 2. Significant differences between day1 and day2 are marked with asterisks (*: *p* < 0.05, **: *p* < 0.01). Curves and bar graphs are presented as mean ± standard error across subjects.

### Activation of brain regions responsive to errors

Regional patterns of activation obtained during the adaptation sessions are reported in Figure 3. For the sake of providing an overall picture of what happened at the brain level during adaptation, we first reported brain activation during the adaptation trials regardless of whether activation was modulated by error magnitude (Figure 3A). Activated regions on day1 and day2 included, as expected for a right-hand movement, the sensorimotor parcels (lobules V, VI, and VIIIb) of the right cerebellum, the left sensorimotor cortical areas (primary motor cortex – BA4, primary somatosensory cortex – BA1, dorsal premotor cortex – BA6, supplementary motor area – BA6), and regions of the left posterior parietal cortex (superior parietal lobule – BA5 and BA7, and extending a bit into the supramarginal gyrus – BA40). Activation foci were also found in the right dorsal premotor cortex, the right superior parietal lobule, and the left sensorimotor cerebellum (lobules V, VI, and VI-IIb). Given the visuomotor nature of the task, activation was also observed in the right and left primary and secondary visual (BA17 and BA18, respectively) cortices. Finally, subcortical regions were also part of the activation network, including the right and left basal ganglia (putamen) and thalamus. There was no activation difference between days 1 and 2.

**Figure 3.**
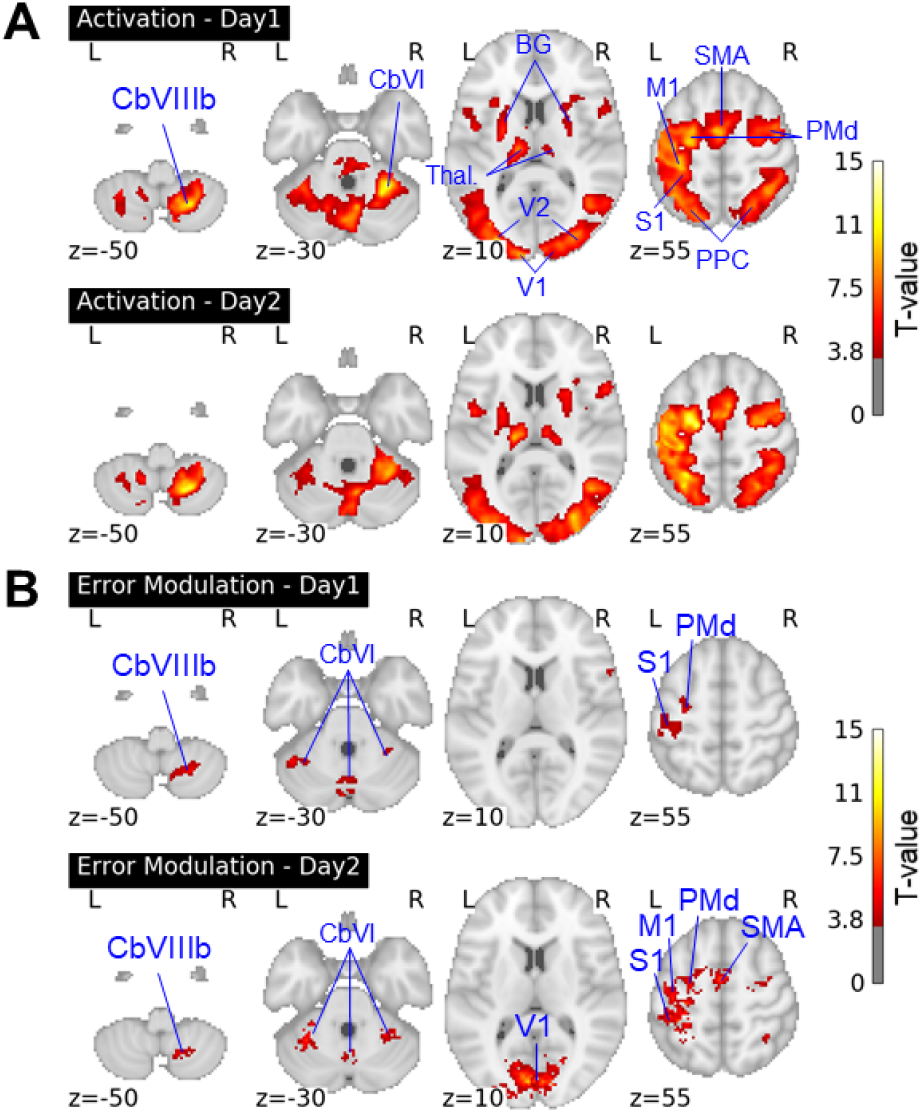
Brain activation. (**A**) evoked by the adaption task, and (**B**) positively related to the magnitude of movement error. M1: primary motor cortex; S1: primary somatosensory cortex; SMA: supplementary motor area; PMd: dorsal premotor cortex; PPC: posterior parietal cortex; BG: basal ganglia; Thal.: thalamus; V1: primary visual cortex; V2: secondary visual cortex; CbVI: cerebellar lobule VI; CbVIIIb: cerebellar lobule VIIIb. R = right hemisphere; L = left hemisphere. x, y, z: coordinates in MNI space.

A subset of regions listed above demonstrated a positive parametric modulation of activation by error magnitude on days 1 and 2 (Figure 3B). Activation foci were located in the right lobule VIIIb and the right and left lobule VI of the cerebellum, in the primary visual cortex, and in the left primary somatosensory and dorsal premotor cortices. Activation was also related to errors in the primary motor cortex and the supplementary motor area on day2. Comparison between days did not reveal any difference in error modulation.

### Connectivity between brain regions responsive to errors

The co-activation patterns derived from the ROIs looked roughly similar and delineated the same error-related network of interconnected regions that spread over the cerebral cortex, the cerebellum, the basal ganglia and the thalamus (Figures 4 & S2). Specifically, they were composed of sensorimotor (BA1, 4, 6), posterior parietal (BA5, 7, 40) and visual (BA17, 18) brain regions, sensorimotor cerebellar territories (lobules V, VI and VIIIb), as well as basal ganglia and thalamus in both the left and right hemispheres. Interestingly, there were also areas located more anteriorly in the frontal lobe, especially in the anterior cingulate cortex (BA 24, 32) and in the lateral prefrontal cortex (BA9, 44). Importantly, the functional connectivity was stronger, on day2 compared to day1, between (i) left S1 and anterior cingulate cortex (MNI coord. = [2, 6, 34], n voxels = 39, pFWE-corr = 0.05; Figures 4A & 5A), (ii) right cerebellar lobule VI and left thalamus (MNI coord. = [-18, -10, 2], n voxels = 49, pFWE-corr = 0.04; Figures 4B & 5A), (iii) right cerebellar lobule VI and right thalamus (MNI coord. = [10, -22, 2], n voxels = 35, marginal effect with pFWE-corr = 0.161 but qFDR-corr = 0.044; Figures 4B & 5A), and (iv) right cerebellar lobule VIIIb and left supramarginal gyrus (MNI coord. = [-48, -30, 42], n voxels = 38, pFWE-corr = 0.031; Figure 4C & 5A).

**Figure 4.**
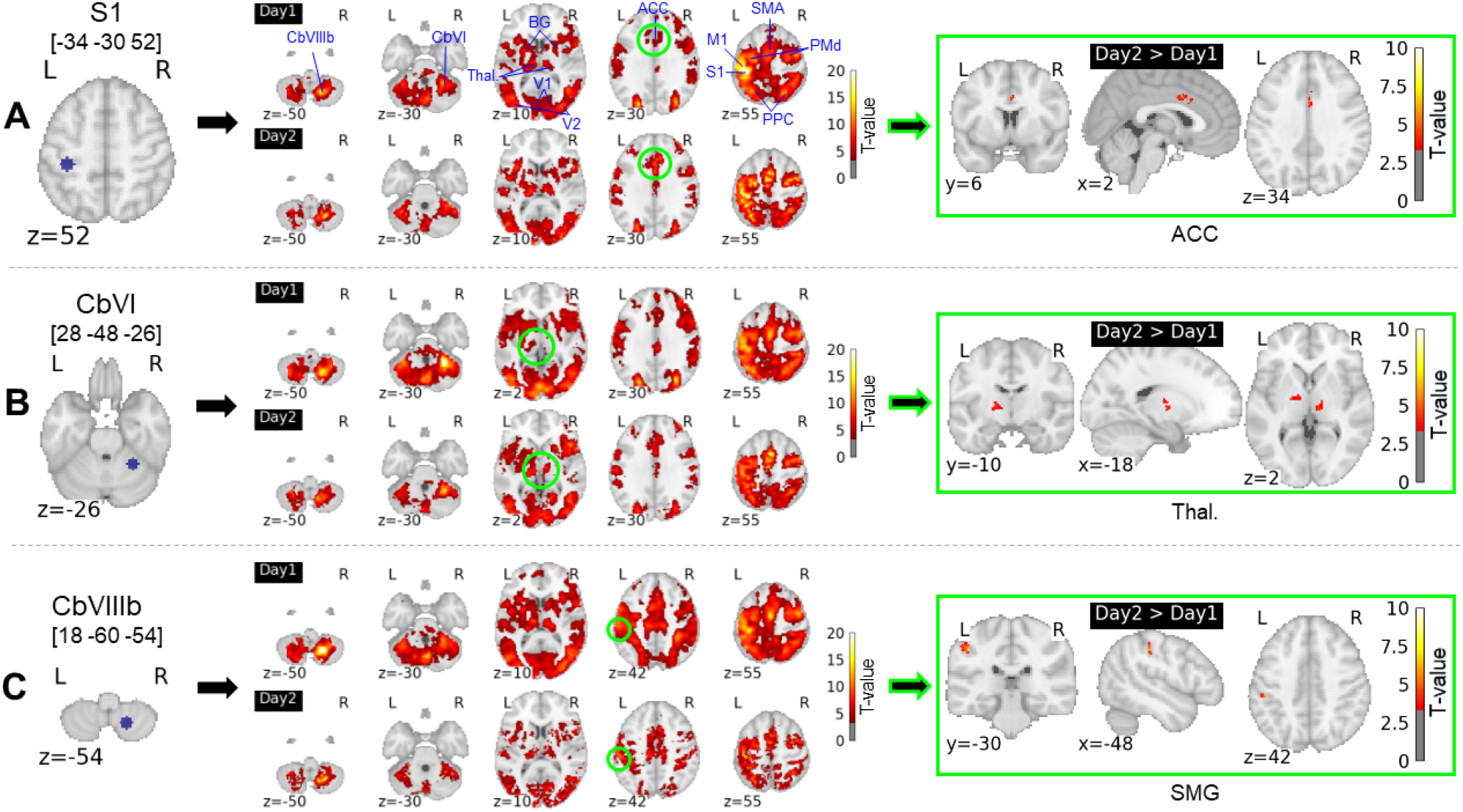
ROI-to-voxel co-activation patterns. The contrast day2>day1 reveals regions whose connectivity with ROIs was stronger on day2 compared to day1. These regions are situated on the connectivity maps using green open circles. ACC: anterior cingulate cortex; M1: primary motor cortex; S1: primary somatosensory cortex; SMA: supplementary motor area; PMd: dorsal premotor cortex; PPC: posterior parietal cortex; SMG: supramarginal gyrus; BG: basal ganglia; Thal.: thalamus; V1: primary visual cortex; V2: secondary visual cortex; CbVI: cerebellar lobule VI; CbVIIIb: cerebellar lobule VIIIb. R = right hemisphere; L = left hemisphere. x, y, z: coordinates in MNI space.

Regarding pathways evoked above between left S1 and anterior cingulate cortex and right cerebellar lobule VI and left thalamus, the increased functional connectivity from day1 to day2 positively correlated with the increased learning rate across individuals (*r* = 0.64, *p* = 0.0007 and *r* = 0.46, *p* = 0.02, respectively; Figure 5B), showing that there was a relationship between increased functional connectivity and savings. Note that these two correlations remained significant when adjusting FDR (4 comparisons) using the Benjamini-Hochberg procedure. On the other hand, regarding the other two pathways between right cerebellar lobule VI and right thalamus and right cerebellar lobule VIIIb and left supramarginal gyrus, there was only a tendency for a positive correlation between increased connectivity and increased learning rate (*r* = 0.34, *p* = 0.1 and *r* = 0.37, *p* = 0.07, respectively; Figure 5B).

**Figure 5.**
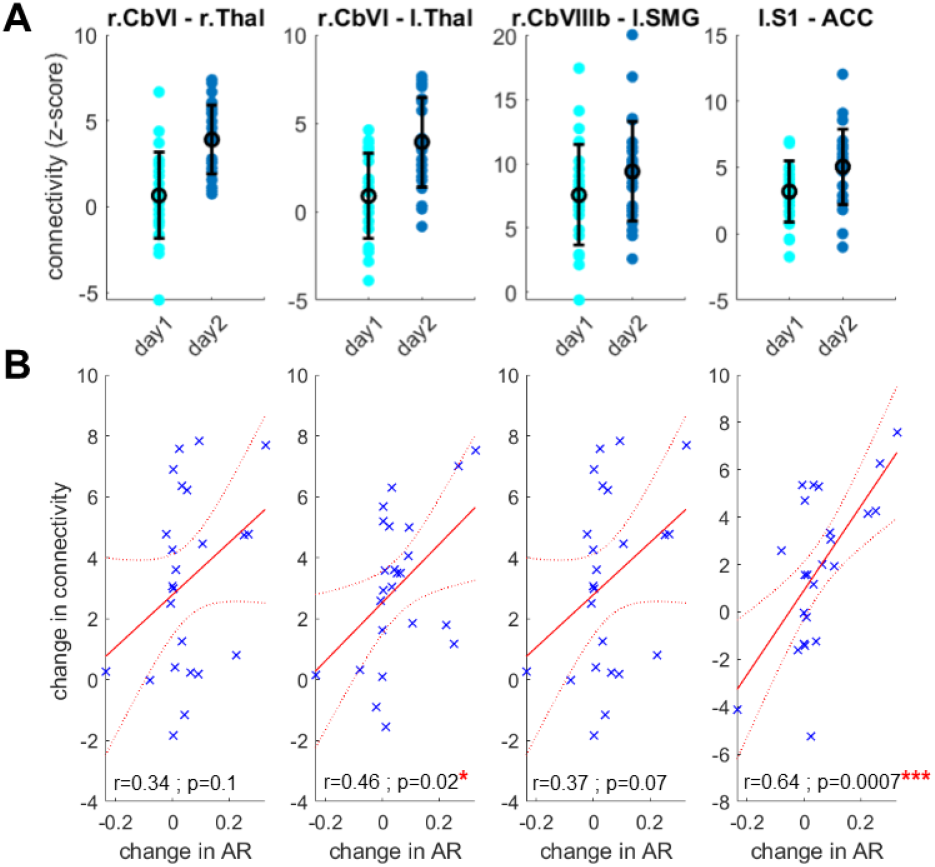
(**A**) Change in interregional co-activation from day1 to day2, and (**B**) its relationship with savings. Savings is represented as change in adaptation rate (AR). ACC: anterior cingulate cortex; S1: primary somatosensory cortex; SMG: supramarginal gyrus; Thal: thalamus; CbVI: cerebellar lobule VI; CbVIIIb: cerebellar lobule VIIIb. r = right hemisphere; l = left hemisphere.

### Changes in intrinsic functional connectivity

ROI-to-voxels analysis did not reveal any difference (F-test) of intrinsic connectivity between the resting state conditions (i.e., pre-day1, post-day1, pre-day2, post-day2). ICA-based analysis revealed a “sensorimotor” network that spread over the supplementary motor area, the anterior (mid) cingulate cortex, the primary sensorimotor cortex, the posterior parietal cortex (BA5 and BA7), the secondary somatosensory cortex and cerebellar regions (Figure 6). This corresponded closely to the network seen in adaptation. Connectivity within this network showed some changes between the resting state conditions, including an increased voxel-to-voxel connectivity following adaptation in both day1 (pre-day1 vs post-day1) and day2 (pre-day2 vs post-day2) in two clusters located centrally in the brain, one in front of the central sulcus (MNI coord. = [-6, -18, 74], n voxels = 167, pFWE-corr = 0.0003) and that covered parts of the primary motor cortex and the dorsal premotor cortex, and the other behind the central sulcus (MNI coord. = [8, -44, 62], n voxels = 290, pFWE-corr = 0.000002) and that covered parts of the primary somatosensory cortex and the posterior parietal cortex (BA5 and BA7).

**Figure 6.**
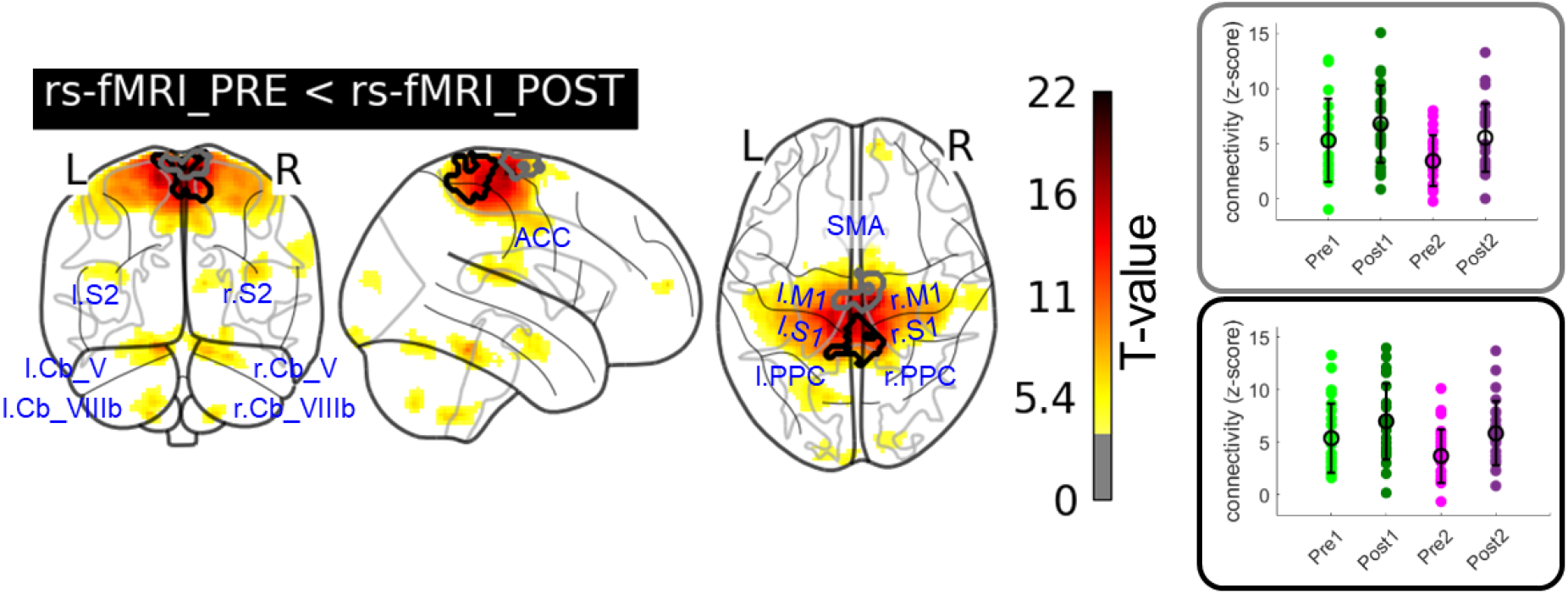
Sensorimotor ICA map obtained during the conditions of resting-state. Clusters that showed increased connectivity following adaptation in both day1 and day2 are represented with black and grey contours. M1: primary motor cortex; S1: primary somatosensory cortex; SMA: supplementary motor area; PPC: posterior parietal cortex; ACC: anterior cingulate cortex; Cb_V: cerebellar lobule V; Cb_VIIIb: cerebellar lobule VIIIb; r.: right hemisphere; l.: left hemisphere.

## Discussion

The present study showed that the more savings there was in a visuomotor learning task, the more important was the increase of connectivity between regions of a cerebello-thalamo-cortical network responsive to errors. This indicated that faster re-adaptation to a perturbation goes through a mechanism that strengthens the communication between brain regions processing errors. Previous studies introduced the notion of memory of errors in which the brain recognizes past errors and commands to increase the gain of error correction, the so-called sensitivity to errors, when re-visiting these errors (Leow et al., 2020; Coltman et al., 2019; Leow et al., 2016; Herzfeld et al., 2014b; Mawase et al., 2014). The present finding suggests that this memory of errors may materialize as a set of brain regions more responsive to errors when re-experiencing a known perturbation.

Specifically, we found that the amount of savings was associated with an increase of connectivity in two pathways, namely a pathway between the primary somatosensory cortex and the dorsal anterior cingulate cortex, and a pathway between the sensorimotor cerebellum (lobule VI) and the thalamus. All these regions have in common that they relate actions to their outcomes, and show error signals when the expected outcomes are not reached. The dorsal anterior cingulate cortex (Klein-Flügge et al., 2022; Hyman et al., 2017; Amiez et al., 2005) and the thalamus (Collomb-Clerc et al., 2023; Chase et al., 2015) are implicated in encoding reward prediction errors, that is the difference between the predicted value of future rewards and their realized value. The primary somatosensory cortex integrates information from both the primary motor cortex and the periphery, making it able to represent discrepancies between expected and actual sensory consequences of the action (Umeda et al., 2019; Mathis et al., 2017). Likewise, the cerebellum represents sensory prediction errors during voluntary movements (Hull, 2020; Brooks et al., 2015; Izawa et al., 2012; Schlerf et al., 2012; Tseng et al., 2007). In the context of visuomotor adaptation, previous observation showed that the error can be broken down into a variety of error types, including reward and sensory errors (Morehead and Orban De Xivry, 2021; Izawa and Shadmehr, 2011). Presumably, the deviation of the visual feedback cursor during pointing movements made erroneous the prediction of the internal model about the sensory consequences of the motor commands (i.e., a sensory prediction error) as well as the expectation about the intrinsic reward associated with successfully attaining the target (i.e., a reward prediction error). With regard to the latter aspect, it is indeed known that the attainment of intrinsic rewards is a general feature of successful completion of goal-directed sensorimotor tasks (Carroll et al., 2019).

The present result relating savings to cerebellar-thalamic connectivity echoes a previous study on locomotor adaptation in which the amount of savings could be predicted from the strength of the cerebellar-thalamic connectivity at rest (Mawase et al., 2017). It was also demonstrated that patients with essential tremor treated with deep brain stimulation of the ventral intermediate nucleus (vim) of the thalamus or vim thalamotomy, which disrupts thalamus functioning, are impaired in their ability to learn from errors (Chen et al., 2006). Furthermore, in fMRI, an increased activity of the cerebellum (lobule VI) correlating with the amount of savings was observed when repeating a motor adaptation task (Debas et al., 2010). These results and the present ones suggest that the rate at which individuals learn from errors is set by the interactions between the cerebellum and the thalamus. It is perhaps surprising, however, that our result did not uncover the entire cerebellothalamo-cortical pathway. Indeed, it is generally assumed that the prediction error generated in the cerebellum is sent to the contralateral thalamus through efferent connections in the superior cerebellar peduncle (Sokolov et al., 2017), which in turn projects the error to multiple cortical targets to adjust the motor command (Abram et al., 2022). Accordingly, there is a possibility that the two pathways reported in the present study are the most visible constituents of a larger-scale network in charge of adjusting the learning rate. We can also mention the fact that single trial activity estimation used in beta series correlation can be quite noisy, which could have affected the sensitivity to detect context-modulated functional connectivity, possibly leaving out some connections. However, there is really no better alternative to assess context-dependent functional connectivity. Results obtained with other methods are roughly similar to those obtained with the beta series method, each method having its pros and cons (Abdulrahman and Henson, 2016; Cisler et al., 2014; Mumford et al., 2012).

Another significant finding was the relationship observed between savings and connectivity of the primary somatosensory cortex. There is still debate about which regions contribute to memory consolidation in motor learning. Some studies found that the primary motor cortex is involved in the retention of newly learned movements (Galea et al., 2011; Orban De Xivry et al., 2011; Hadipour-Niktarash et al., 2007; Muellbacher et al., 2002), while others did not find it to be involved (Kumar et al., 2019; Baraduc et al., 2004). On the other hand, several studies supported the view that the primary somatosensory cortex, instead of the primary motor cortex, is key to retention (Ebrahimi and Ostry, 2024; Darainy et al., 2023; Kumar et al., 2019). According to these studies, this cortex would store learning-updated sensory states which serve to guide the movement. The present results support a participation of the primary somatosensory cortex to memory consolidation, as reflected by a faster relearning. However, the fact that savings was associated with multiple connectivity pathways suggests that memory consolidation is not limited to a given brain region but rather depends on interregional interactions.

One important issue in motor adaptation has been to understand which processes contribute to error-based learning. Behavioural studies revealed a dual-process organization with a slow/implicit learning process and a fast/explicit learning process (Huberdeau et al., 2015). However, their respective contributions to savings vary across studies, including an improvement of the slow/implicit process (Yin and Wei, 2020; Joiner and Smith, 2008; Smith et al., 2006), of the fast/explicit process (Avraham et al., 2021; Morehead et al., 2015), and of both (Coltman et al., 2019). This variety of findings comes from the fact that the interplay between the two processes critically depends on the specifics of the learning protocol. In a standard protocol of visuomotor adaptation, as the present one, both processes are involved, and as such may have contributed to savings. This assumption is supported by the contribution of cerebellar and cortical regions in the connectivity pathways that were related to savings. Indeed, the slow/implicit and fast/explicit components of savings would be implemented in the cerebellum and the cerebral cortex, respectively (Kim et al., 2015). Future research on the neural correlates of savings should consider using more sophisticated protocols able to isolate the specific contributions of these two processes of adaptation (Morehead et al., 2017; Haith et al., 2015; Taylor et al., 2014).

Finally, following each session of adaptation, there was evidence of an increased connectivity within regions of the resting-state sensorimotor network (i.e., somatosensory cortex and frontal motor areas). This observation is consistent with the idea that motor adaptation tasks drive plasticity in both sensory and motor brain areas, plasticity in the sensory and motor systems being reciprocally linked (Ostry and Gribble, 2016; Vahdat et al., 2011). However, we failed to establish a relationship between these changes in the restingstate sensorimotor network and the (SSM) learning parameters, which raises the question of whether these changes were specific to the adaptation task or reflected transient changes that would occur following any task execution. A recent study which investigated changes in the resting-state connectivity following constant perturbation (i.e., adaptation) and random perturbation (i.e., no adaptation) tasks in a force field environment linked most changes in intrinsic connectivity to the adaptation process (Farrens et al., 2023). Accordingly, it seems reasonable to assume that the observed changes in the resting state sensorimotor network were driven by the adaptation process. Finally, we found that intrinsic connectivity of the sensorimotor network was the same following both sessions of adaptation (i.e., no change accumulated from session 1 to session2). Thus, the changes reported in task-evoked (error-related) responses of the brain during the second session of adaptation did not seem to rely on a change in intrinsic brain connectivity. This null finding is rather surprising in view of the role of intrinsic brain activity in shaping task activations (Cole et al., 2016; Tavor et al., 2016; Cole et al., 2014) and organizing brain functions (Raichle, 2015). Studies with more intensive learning (e.g., more movement trials, multiple sessions), which should magnify behavioural and connectivity changes, may be valuable to revisit this plasticity issue and investigate the relationship between intrinsic and task-evoked activity in motor adaptation.

In summary, we examined which changes happened in the brain of individuals who showed savings when relearning a visuomotor rotation task. We found an increased functional connectivity between subcortical and cortical regions responsive to movement errors during relearning, which was predictive of the amount of savings. Hence, when individuals better perform a motor task than before it is because of an improved communication between brain regions involved in error processing. This provides a concrete brain mechanism to the concept of increased sensitivity to errors used in previous studies on motor learning and savings. However, such an improved communication did not persist in the resting brain following learning of the motor task.

## Acknowledgements

This work was performed on the IRMaGe platform member of France Life Imaging network (grant ANR-11-INBS-0006) and was supported by NeuroCoG IDEX UGA, “Investissements d’avenir” program (grant ANR-15-IDEX-02). We would also like to thank Marie Latil for her help in resting-state fMRI analysis as well as Johan Pietras and Lucie Miquel for their contribution in MRI data collection.

## Authors contributions

Conceptualization: LS, RL, TI, VN, DJO, FC ; Methodology: LS, LL, PAB, FC; Software: LS, LL, FC; Formal analysis: LS, FC; Investigation: LS, AC, FC; Data curation: LS, FC; Visualization: LS, FC; Writing - original draft: LS; Writing – review & editing: LL, PAB, AC, RL, TI, VN, DJO, FC; Supervision: FC; Funding acquisition: FC.

## Supplementary Information

**Figure S1.**
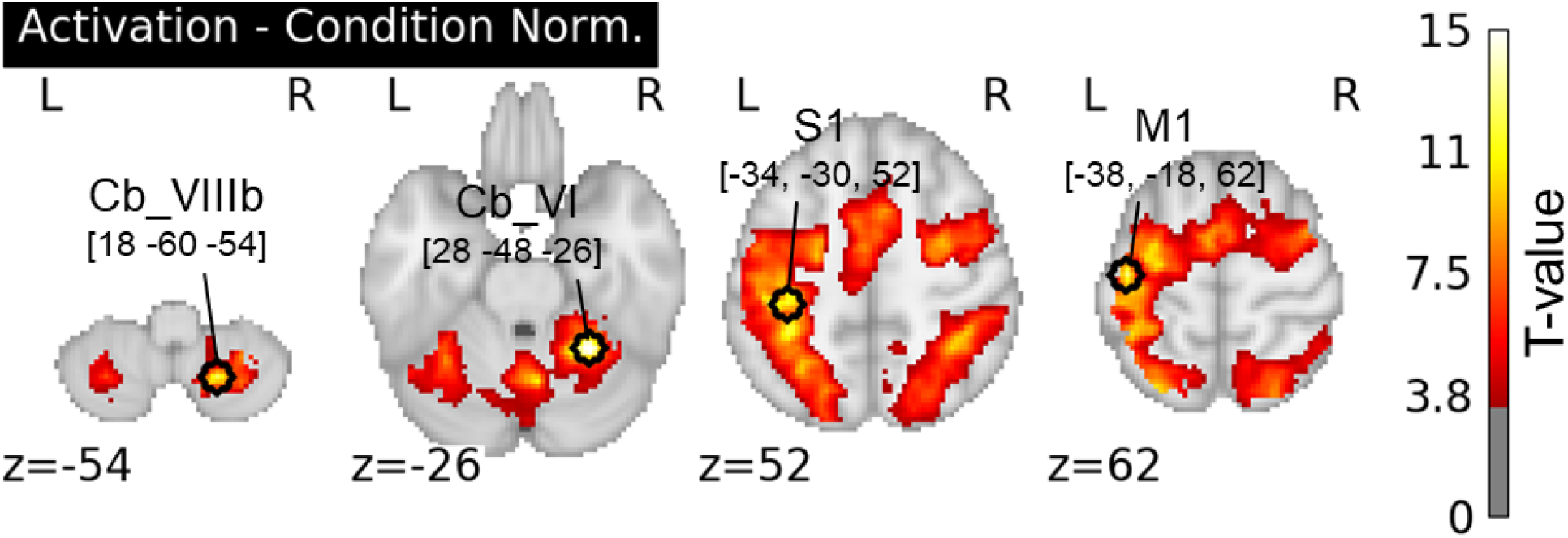
ROIs locations. ROIs have been defined from local maxima of the group-level activation map of the Norm condition (against implicit baseline). M1: primary motor cortex; S1: primary somatosensory cortex; Cb_VI: cerebellar lobule VI; Cb_VIIIb: cerebellar lobule VIIIb; L: left hemisphere; R: right hemisphere.

**Figure S2.**
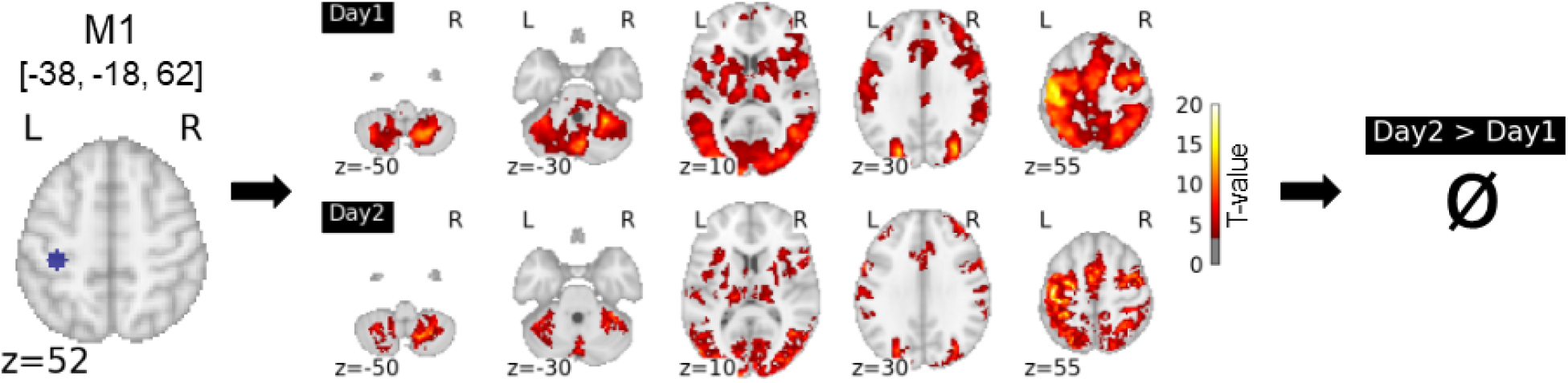
M1-to-voxels co-activation (or equivalently functional connectivity) patterns between regions responsive to errors. The contrast day2>day1 indicated that M1 connectivity related to error did not change on day2 compared to day1.

